# The involvement of the ER-phagy receptor FAM134B in membrane contact sites between ER and endolysosomes promotes ERLAD

**DOI:** 10.1101/2025.07.08.663682

**Authors:** Elisa Fasana, Ilaria Fregno, Maurizio Molinari

## Abstract

Membrane contact sites (MCS) between organelles maintain the proximity required for controlled exchange of small molecules and ions yet preventing fusion events that would compromise organelles’ identity and integrity. Here, by investigating the intracellular fate of the disease-causing Z-variant of alpha1 antitrypsin (ATZ), we report on a novel function of MCS between the endoplasmic reticulum (ER) and RAB7/LAMP1-positive endolysosomes in ER-to-lysosome-associated degradation (ERLAD). For this function, the VAPA:ORP1L:RAB7 multi-protein complex forming MCS between the ER and endolysosomes engages, in an ERLAD client-driven manner, the misfolded protein segregation complex formed by the lectin chaperone Calnexin (CNX), the ER-phagy receptor FAM134B and the ubiquitin-like protein LC3. Generation of this supramolecular complex facilitates the membrane fusion events regulated by the SNARE proteins STX17 and VAMP8 that ensure efficient delivery of ATZ polymers from their site of generation, the ER, to the site of their intracellular clearance, the degradative RAB7/LAMP1-positive endolysosomes.

## Introduction

Proteins that fail to attain the native structure in the ER are retro-translocated into the cytosol for proteasomal clearance by ERAD pathways that have been studied in great detail in the last 35 years^1-3^. An increasing number of disease-causing misfolded proteins that cannot engage the ERAD machinery have been described in recent years. Often, these are large or aggregated polypeptides that fail to be retro-translocated across the ER membrane. Rather, they are segregated in ER subdomains that eventually deliver their toxic content to endolysosomal degradative compartments via ERLAD, an acronym that we introduced to define all autophagic and non-autophagic pathways that ensure the lysosomal clearance of ERAD-resistant misfolded polypeptides from the ER^4-8^. ATZ is a model ERLAD client^9-12^. An aspartic acid to lysine (E342K) point mutation enhances polymerogenicity of this disease-causing polypeptide^13^. ATZ polymers fail dislocation across the ER membrane for ERAD. They undergo repeated cycles of de-glucosylation and re-glucosylation of the ATZ N-glycan at position 83 by ER-resident α-glucosidase II and UDP-glucose:glycoprotein glucosyltransferase 1 (UGT1), which prolong the association of ATZ polymers with the lectin chaperone CNX^9^. ATZ:CNX complexes eventually engage the ER-phagy membrane receptor FAM134B and lipidated LC3 to favor segregation of polymeric ATZ in ER subdomains and/or in ER-derived vesicles that release their toxic content within RAB7/LAMP1-positive endolysosomes upon STX17:VAMP8-driven fusion of ER and endolysosomal membranes^9,10^. How ER subdomains and/or ER-derived vesicles containing ATZ polymers are docking to the appropriate degradative compartment to deliver their toxic content for clearance is not known.

Membrane contact sites (MCS) are sites of proximity between the membrane of organelles. They are stabilized by tethering machineries that maintain organelles’ proximity but prevent membrane fusion events that would compromise organelle identity and integrity^14,15^. The MCS between ER and endolysosomes are well-characterized, are maintained by VAPA-ORP1L-RAB7 multiprotein complexes, and serve as site of lipids and calcium transfer^14-21^.

The finding that VAPA is a major interactor of FAM134B^22^, as well as transmission electron microscopy (TEM) micrographs showing the proximity of ER subdomains containing ATZ polymers and endolysosomes before and during delivery of the ERLAD client within the degradative compartment (**Fig. 1**)^10^, led us to examine if the generation of contact sites is required for lysosomal protein clearance from the ER. Here, we show that cells producing ATZ polymers, i.e., misfolded polypeptides to be selected for lysosomal clearance (i.e., ERLAD), display higher number of ER-endolysosomes MCS compared to cells that do not express misfolded proteins, or that express the NHK variant of alpha1 antitrypsin, a folding-defective polypeptide that is efficiently retro-translocated across the ER membrane for proteasomal degradation (ERAD)^12,23-25^. We report that the *segregation complex* formed at the ER membrane by the lectin chaperone CNX and the ER-phagy receptor FAM134B and stabilized upon intralumenal expression of ATZ polymers^10^ also contains the ER membrane protein VAPA, the cytoplasmic bridging protein ORP1L and the small RAB7 GTPase at the limiting membrane of the endolysosomes. This generates ER-endolysosome MCS that facilitate delivery of the ERLAD client within degradative compartments upon membrane fusion events controlled by the SNARE proteins STX17 and VAMP8. Inhibiting the formation of ER-endolysosomes MCS upon genome editing to delete the VAPA or the ORP1L proteins substantially hampers lysosomal clearance of misfolded ATZ.

**Figure 1.**
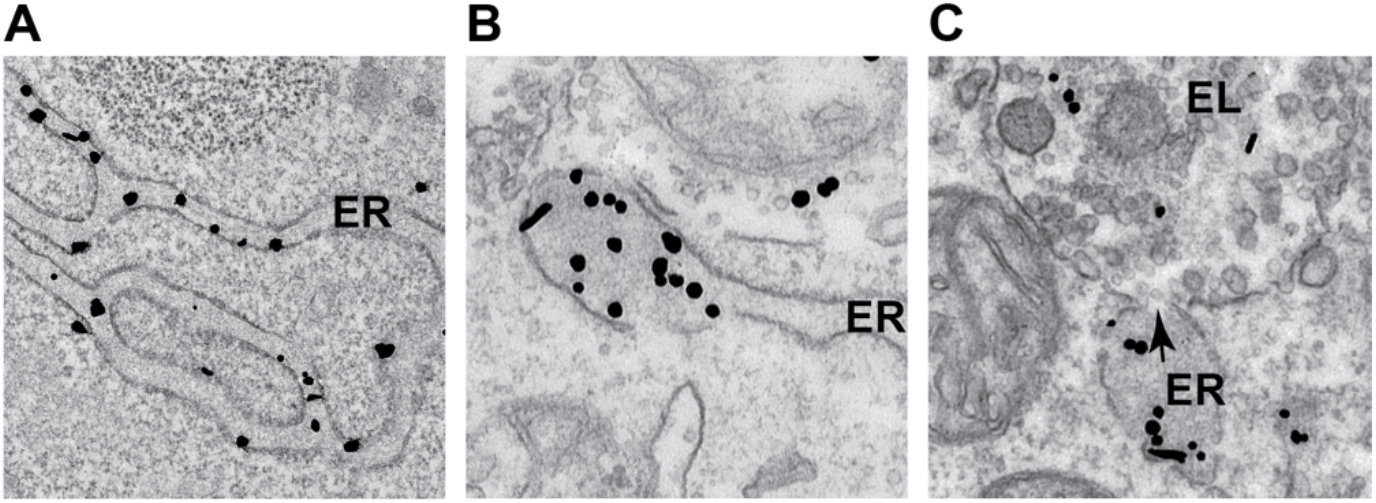
Segregation and lysosomal delivery of ATZ polymers. (**A**) RT-TEM micrograph showing immunogold-labeled ATZ polymers dispersed in the endoplasmic reticulum (ER) lumen. (**B**) ATZ polymers are segregated in terminal portions of the ER. (**C**) The arrow shows the site, where the membrane of the ER subdomain fuses with the membrane of the endolysosomal degradative compartment (EL).

## Results

### ATZ is delivered from the ER lumen into endolysosomes for clearance

Room Temperature (RT)-TEM has been used to monitor the fate of disease-causing ATZ polymers generated in the ER lumen (Fig. 1A, ER), and segregated in ER subdomains (Fig. 1B) that eventually fuse with endolysosomes (EL) to deliver their toxic content for clearance (arrow in Fig. 1C)^9,10^.

Inactivation of lysosomal hydrolases upon exposure of mouse embryonic fibroblasts (MEF) to Bafilomycin A1 (BafA1) results in the accumulation of undegraded ATZ polymers that have been delivered within RAB7/LAMP1-positive endolysosomes (confocal laser scanning microscopy (CLSM), Fig. 2A and quantifications with LysoQuant^26^, a deep-learning-based analysis sojware for segmentakon and classificakon of fluorescence images, Fig. 2E)^10^. Lysosomal clearance of ATZ requires the contact between ER-derived structures containing ATZ polymers (ER in Fig. 1C, highlighted in red in the immunoelectron microscopy (IEM) micrograph in Fig. 2B) and the membrane of degradative endolysosomes (EL in Fig. 1C, highlighted in green in Fig. 2B). The contact between ER and endolysosome is followed by a membrane fusion event involving the SNARE proteins Syntaxin17 (STX17) and VAMP8 that we have previously characterized^10^, and that delivers the ATZ polymers from the ER lumen into the lumen of the RAB7/LAMP1-positive degradative endolysosomes (the arrow in Fig. 1C and the blue asterisk in Fig. 2B show the opening between the ER and the EL membranes). In cells lacking STX17 generated by CRISPR/Cas9 genome editing^10^, the delivery of ATZ within the LAMP1-positive degradative compartments is strongly inhibited (Figs. 2C, 2E)^10^. The ER structures segregating ATZ polymers (Fig. 2D, highlighted in red) dock at the limiting membrane of the RAB7/LAMP1-positive endolysosomes (highlighted in green) but fail to fuse and to deliver ATZ polymers in cells with dysfunctional SNARE complexes (Figs. 2C-2E).

**Figure 2.**
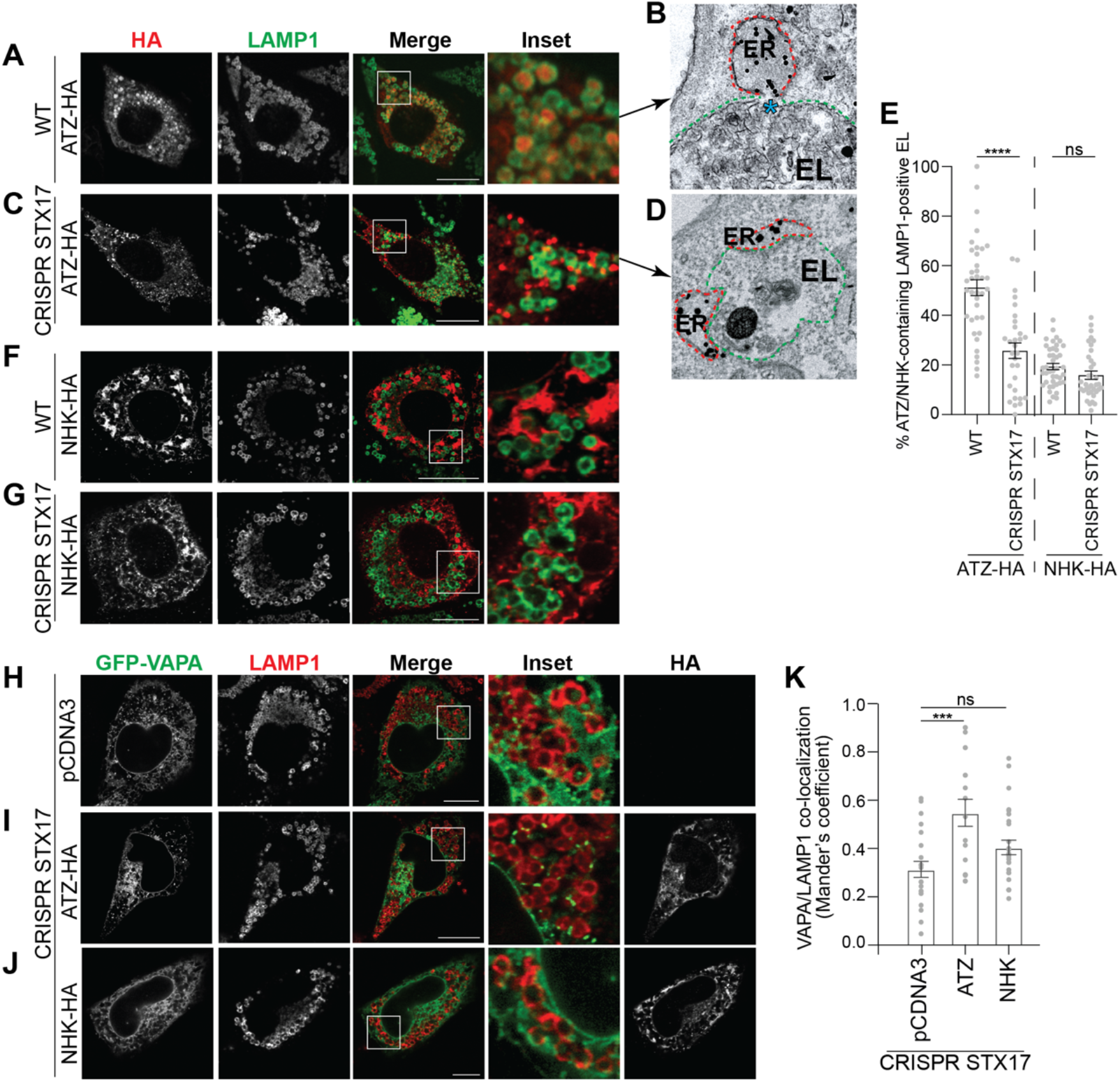
ATZ expression increases ER-endolysosome contact sites. (**A**) CLSM analyses of ATZ-HA delivery (and accumulation) within LAMP1-positive endolysosomes in wild type MEF treated with 50nM BafA1. (**B**) Same as **A** in RT-IEM. The blue asterisk shows the site of fusion between ATZ-enriched ER subdomains (dotted red line) and the degradative organelle (dotted green line). (**C**) Accumulation of ER containing ATZ-HA in proximity of LAMP1-positive endolysosomes in CRISPR STX17 MEF treated with 50nM BafA1. (**D**) Same as **C** in RT-IEM. (**E**) LysoQuant quantification of ATZ delivery within LAMP1-positive endolysosomes in **A, C, F, G** (n = 37, 30, 42 and 36 cells of 3 independent experiments, respectively). Unpaired two-tailed *t*-test, ^ns^*P* > 0.05, ****P < 0.0001. t=5.605 and t=1.686, respectively. (**F**) NHK-HA does not accumulate within LAMP1-positive endolysosomes in wild type MEF treated with 50nM BafA1. (**G**) NHK-HA distribution in CRISPR STX17 MEF treated with 50nM BafA1. (**H**) CLSM analysis showing GFP-VAPA and LAMP1 distribution in CRISPR STX17 MEFs mock-transfected, (**I**) expressing ATZ-HA, or (**J**) expressing NHK-HA. (**K**) Mander’s coefficient of VAPA/LAMP1 co-localization of **H-J** (n= 22, 19 and 25 cells of 3 independent experiments, respectively). One-way ANOVA and Dunnett’s multiple comparisons test, ^ns^*P* > 0.05, \*\*\**P* < 0.001. F=5.825. Each data point shows the level of co-localization between VAPA and LAMP1. 1.0, complete juxtaposition of the two channels. 0.0, no juxtaposition. Scale bar: 10μm.

NHK, a folding-defective variant of alpha1-antitrypsin that is mainly degraded by the ubiquitin/proteasome system via ERAD^12,23-25^ is only poorly delivered within the endolysosomal compartment in MEF (Figs. 2F, 2E) and deletion of STX17 does not affect its intracellular distribution (Figs. 2G, 2E)^12^.

### Luminal accumulation of ATZ promotes formation of ER-endolysosomes contact sites

Next, we checked by CLSM whether the expression of the ERLAD client ATZ, or of the ERAD client NHK affects the proximity between the ER and the endolysosomes. The experiments were performed in CRISPR STX17 cells, where the absence of the fusion machinery that drives the transfer of ATZ polymers from the ER to the lumen of endolysosomes eventually stabilizes the juxtaposition of the two organelles^10^. The Mander’s correlation coefficient used to establish the level of co-localization between the ER marker protein VAPA and the endolysosomal marker protein LAMP1^27-29^ reveals higher values in cells expressing ATZ than in cells mock-transfected or expressing the folding-defective ERAD client NHK (Figs. 2H-2K, respectively). Thus, the expression of a misfolded polypeptide that must be cleared by the endolysosomal system enhances the contacts between the biosynthetic ER compartment and the degradative endolysosomal compartments, whereas the expression of a misfolded protein mainly cleared by the ubiquitin/proteasome system does not.

### The intraluminal expression of ATZ enhances the co-localization of contact sites regulators at the membrane of endolysosomes

To confirm that the luminal accumulation of the ERLAD client ATZ promotes the formation of contacts between ER subdomains and endolysosomes, we monitored the subcellular localization of proteins reportedly involved in ER-endolysosomes contact sites, which are set by the VAPA localized at the ER membrane, the small GTPase RAB7 associated at the membrane of LAMP1-positive endolysosomes, and the cytoplasmic protein ORP1L that bridges the two^30-33^. Analyses by CLSM in CRISPR STX17 cells reveal that in cells expressing HALO-ATZ, the misfolded polypeptide (orange arrowheads in the Insets, Fig. 3A) co-localizes with GFP-VAPA (green arrowheads) and with V5-ORP1L (purple arrowheads) at the limiting membranes of LAMP1-positive endolysosomes (white arrowheads). In cells expressing the ERAD client NHK, we observe two major differences that suggest that the ERAD client does not induce VAPA/ORPL1 contact sites formation. First, the lack of condensation of the V5-ORP1L signal in correspondence to the LAMP1 signal identifying the limiting membrane of the endolysosomal compartments (Fig. 3B, compare the subcellular distribution of V5-ORP1L in the Inset, with the corresponding Inset of Fig. 3A).

**Figure 3.**
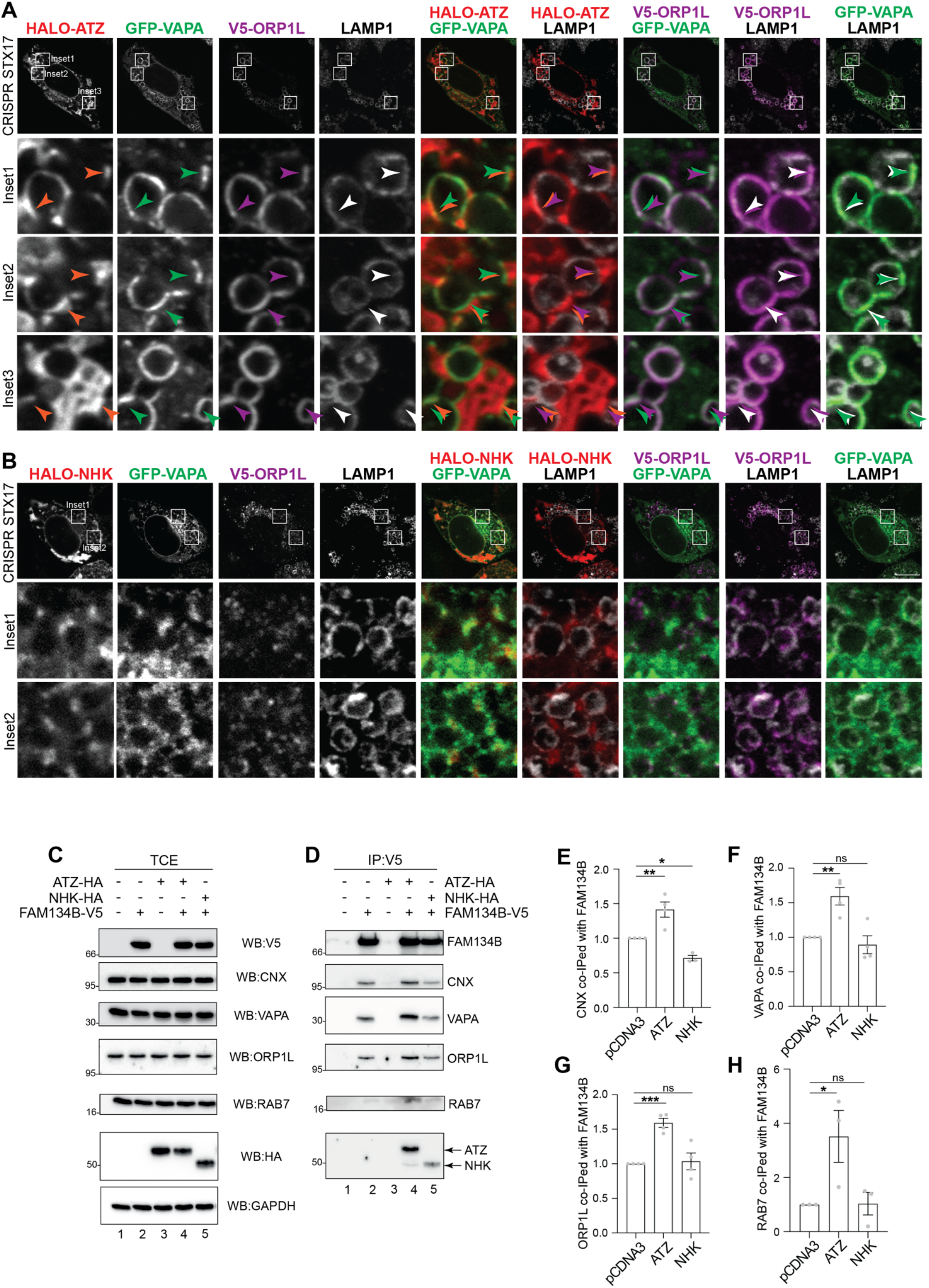
Expression of ATZ induces the formation of CNX:FAM134B:VAPA:ORP1L:RAB7 complexes. (**A**) CLSM analysis showing GFP-VAPA, V5-ORP1L and LAMP1 distribution in HALO-ATZ transfected CRISPR STX17 MEFs treated with 50nM BafA1. (**B**) Same as **A** for cells expressing HALO-NHK. (**C**) WB of the total cell extract (TCE) of HEK293 cells transfected with empty vector (lane 1), FAM134B-V5 (lane 2), ATZ-HA (lane 3), FAM134B-V5 and ATZ-HA (lane 4), or FAM134B-V5 and NHK-HA (lane 5), incubated for 6 h with 100 nM BafA1 and then lysed with 2% CHAPS. Membranes were probed with anti-V5, anti-CNX, anti VAPA, anti-ORP1L, anti-Rab7, anti-HA and anti-GAPDH antibodies, respectively (**D**) WB of the anti-V5 immunocomplexes of **C.** Membranes were probed with anti-V5, anti-CNX, anti-VAPA, anti-ORP1L, anti-Rab7 and anti-HA antibodies, respectively. (**E**) Quantification of the co-immunoprecipitation of CNX with FAM134B observed in **D**, the association in mock-transfected cells is set to 1. One-way ANOVA and Dunnett’s multiple comparisons test, \**P* < 0.1, \*\**P* < 0.01. F=23.06. n=4 independent experiments for pCDNA3 and ATZ, n=3 independent experiments for NHK. (**F**) Quantification of the co-immunoprecipitation of VAPA with FAM134B observed in **D**. One-way ANOVA and Dunnett’s multiple comparisons test, ^ns^*P* > 0.05, \*\**P* < 0.01. F=13.00. n=4 independent experiments (**G**) Quantification of the co-immunoprecipitation of ORP1L with FAM134B observed in **D**. One-way ANOVA and Dunnett’s multiple comparisons test, ^ns^*P* > 0.05, \*\*\**P* < 0.001. F=26.58. n=4 independent experiments (**H**) Quantification of the co-immunoprecipitation of RAB7 with FAM134B observed in **D**. One-way ANOVA and Dunnett’s multiple comparisons test, ^ns^*P* > 0.05, \**P* < 0.1. n=3 independent experiments F=5.540. Scale bar: 10μm.

Second, and consistent with the role of ORP1L in bridging the ER-localized VAPA protein by anchoring RAB7 GTPase located at the endolysosomal membrane^30-33^, GFP-VAPA does not accumulate near the rim of LAMP1 enriched endolysosomes. (Fig. 3B), as a significant difference with cells expressing the ERLAD client ATZ (Fig. 3A).

### The intraluminal expression of ATZ promotes the formation of CNX:FAM134B:VAPA:ORP1L:RAB7 complexes

Few considerations led us to examine a possible induction of protein complexes engaging VAPA, ORP1L and RAB7 that control ER-endolysosomes contact sites in cells accumulating ATZ. First, the increased interaction between proteins involved in the generation/stabilization of ER-endolysosomes contact sites seems a *conditio sine qua non* to explain the imaging data shown in Figs. 2, 3A, 3B testifying the spatial co-localization of these proteins. Second, previous experiments from our lab showed that the luminal accumulation of ATZ relies on the persistent association of the misfolded polypeptide with the ER-resident lectin chaperone CNX via the N-linked glycan at position 83 of ATZ. The long-lasting association promotes the engagement of the ER-phagy receptor FAM134B^9,10^. Third, VAPA is a major interactor of ER-phagy receptor FAM134B^22^. However, the function of the VAPA:FAM134B complexes is unknown. To verify if luminal accumulation of misfolded proteins affects the propensity of ER and endolysosomes to engage in contacts, HEK293 cells were mock-transfected (Fig. 3C, lane 1), or were transfected with plasmids for expression of FAM134B-V5 (lane 2), ATZ-HA (lane 3), FAM134B-V5 and ATZ-HA (lane 4), or FAM134B-V5 and NHK-HA (lane 5). FAM134B was immunoisolated with an anti-V5 antibody from lysates of cells solubilized with the zwitterionic detergent CHAPS to preserve the association of the ER-phagy receptor with ectopically expressed and endogenous proteins in functional complexes^9,10^ (Fig. 3D). In mock-transfected cells, no V5 signal is visible, confirming the specificity of the anti-V5 antibody (Fig. 3D, lane 1). Immunoisolation of ectopically expressed FAM134B confirms the constitutive association of endogenous CNX (lane 2)^10^ and of endogenous VAPA (lane 2)^22^ revealed with the corresponding antibodies in the western blots. Significantly, these complexes also contain endogenous ORP1L and low, but detectable amounts of endogenous RAB7 (lane 2), reporting on the constitutive presence of ER-endolysosomes contact sites^30,32^. The luminal accumulation of the ERLAD client ATZ enhances the formation/stability of the functional complexes between FAM134B and endogenous CNX (Figs. 3D, lanes 2 *vs*. 4, 3E), as we previously reported^10^. Our new tests add that ATZ expression also substantially increases the association of endogenous VAPA, ORP1L, and RAB7 (lanes 2 *vs*. 4, 3F-3H) with the functional FAM134B:CNX complex that ensures lysosomal clearance of ATZ.

NHK, a misfolded polypeptide cleared from cells via ERAD^12,23-25^, only weakly engages FAM134B and the ERLAD machinery (lane 5)^12^ and its luminal expression does not affect the co-precipitation of endogenous CNX, VAPA, ORP1L and RAB7 with FAM134B (Figs. 3D, lanes 2 *vs*. 5 and 3E-3H).

### VAPA is required for ERLAD of ATZ

VAP proteins are present in several flavors in the mammalian ER, VAPA and VAPB being involved in ER-endolysosomes, and ER-mitochondria contact sites, respectively^31,34,35^. To assess the involvement of the VAPA:ORP1L complex in the clearance of misfolded ATZ from the ER, we first monitored by CLSM the lysosomal delivery of ATZ in wild type HeLa cells and in HeLa cells, where VAPA and VAPB have been deleted (Fig. 4A)^36^. The deletion of VAPA/B (Fig. 4B, lower panels and quantification by LysoQuant, 4C) substantially inhibits delivery of ATZ within LAMP1-positive endolysosomes compared to wild type HeLa cells (Fig. 4B, upper panels and 4C). The back-transfection of GFP-VAPA partially restores the lysosomal delivery of ATZ in VAPA/B double KO cells (Figs. 4D, middle panel, 4E). The back-transfection of GFP-VAPB does not restore it (Figs. 4D, lower panel, 4E). These data show that VAPA, and not VAPB, plays a crucial role in the stabilization of ER-endolysosome contact sites that promote ATZ clearance from the ER. Co-immunoprecipitation essays show that in cells lacking VAPA (Figs. 4F, 4G, lanes 5-8) the luminal accumulation of ATZ does not augment the interaction between (i.e., the co-precipitation of) CNX and FAM134B (Figs. 4G, lanes 5-8 and quantification in 4H). The absence of VAPA abolishes the engagement of endogenous ORP1L in the complex that should warrant the formation of the ER-endolysosome contact sites (Figs. 4G, compare lane 4 (co-precipitation of the contact sites components in wild type HeLa cells *vs*. lane 8 the same in HeLa cells lacking the VAPA component, 4J). This confirms, in another cell line, that FAM134B associates with VAPA (as shown in Fig. 3 for MEF)^22,37-42^ and that VAPA is required to engage ORP1L in functional complexes promoting the formation of contacts between the biosynthetic and the degradative compartments.

**Figure 4.**
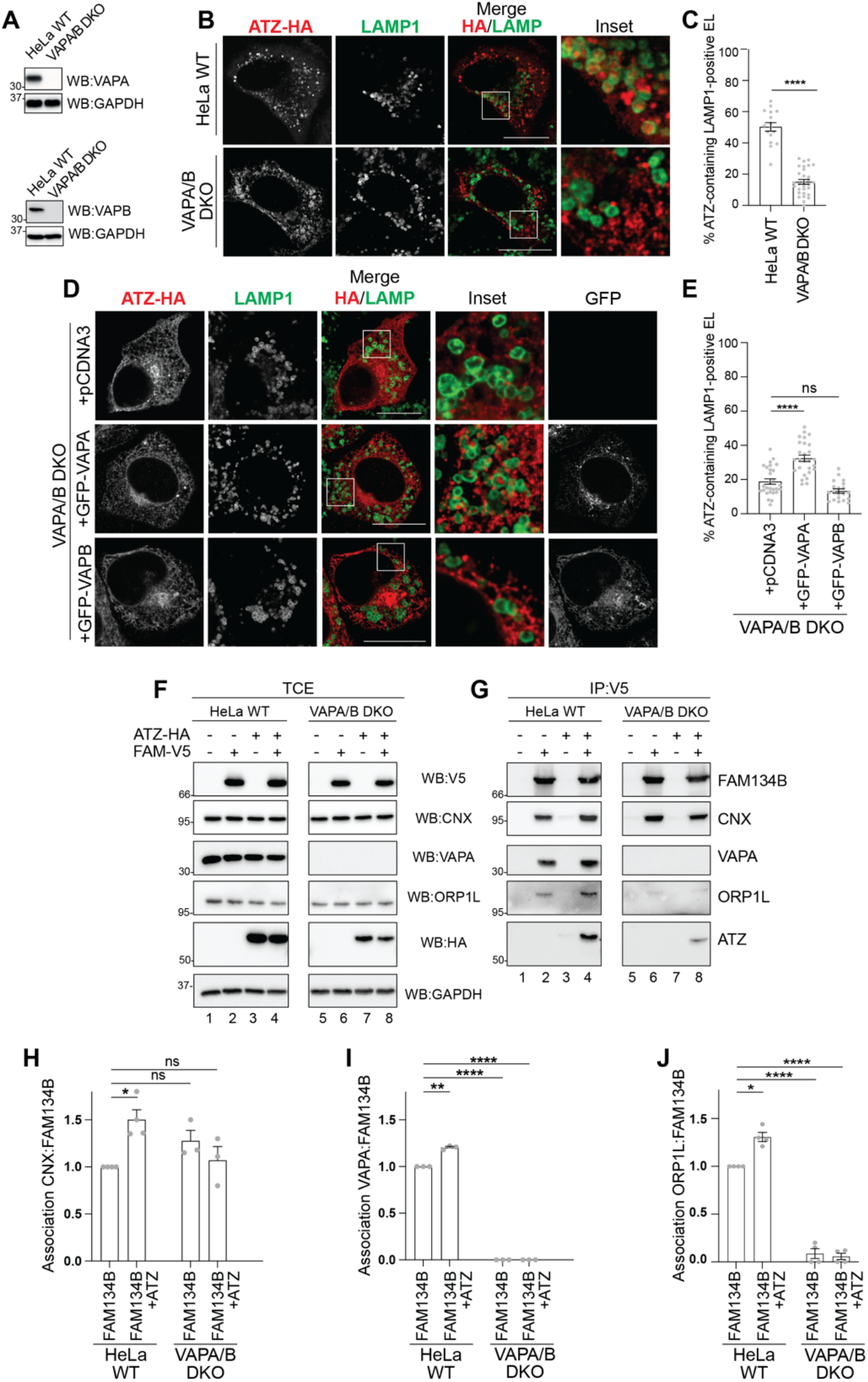
VAPA is required for ATZ delivery to endolysosomes. (**A**) WB analysis showing VAPA (upper) and VAPB (lower panels) down-regulation efficiency for HeLa VAPA/B DKO cells. (**B**) CLSM analysis of ATZ-HA delivery to LAMP1-positive endolysosomes in HeLa WT (upper panels) and in HeLa VAPA/B DKO cells (lower panels) treated with 100nM BafA1. (**C**) LysoQuant quantification of **B** (n= 16 and 30 cells of 3 independent experiments, respectively). Unpaired two-tailed *t*-test, ^ns^*P* > 0.05, ****P < 0.0001. t=12.11. (**D**) CLSM analysis of ATZ-HA delivery to LAMP1-positive endolysosomes in HeLa VAPA/B DKO cells treated with 100nM BafA1 mock transfected (upper panels), transfected with GFP-VAPA (middle panels) or with GFP-VAPB (lower panels). (**E**) Quantification of **D** (n = 28, 27, and 20 cells of 3 independent experiments, respectively). One-way ANOVA and Dunnett’s multiple comparisons test, \*\**P* < 0.01, \*\*\*\**P* < 0.0001. F=34.62. (**F**) WB of the total cell extract (TCE) of HeLa WT cells (lanes 1-4) and HeLa VAPA/B DKO cells (lanes 5-8) transfected with empty vector (lanes 1 and 5), FAM134B-V5 (lanes 2 and 6), ATZ-HA (lanes 3 and 7), or FAM134B-V5 and ATZ-HA (lanes 4 and 8), incubated for 6 h with 100 nM BafA1 and then lysed with 2% CHAPS. Membranes were probed with anti-V5, anti-CNX, anti-VAPA, anti-ORP1L, anti-HA and anti-GAPDH antibodies, respectively. (**G**) WB of anti-V5 immunocomplexes of **F**. Membranes were probed with anti-V5, anti-CNX, anti-VAPA, anti-ORP1L, and anti-HA antibodies, respectively. **H)** Quantification of the association CNX:FAM134B observed in **G**. One-way ANOVA and Dunnett’s multiple comparisons test, ^ns^*P* > 0.05, \**P* < 0.1. F=11.65. n=4 independent experiments for HeLa WT and n=3 for HeLa VAPA/B DKO cells. (**I**) Quantification of the association VAPA:FAM134B observed in **G**. One-way ANOVA and Dunnett’s multiple comparisons test, \*\**P* < 0.01, **** *P* < 0.0001. F=329.2. n=3 independent experiments (**J**) Quantification of the association ORP1L:FAM134B observed in **G**. One-way ANOVA and Dunnett’s multiple comparisons test, \**P* < 0.1, **** *P* < 0.0001. F=148.9. n=4 independent experiments. Scale bar: 10μm.

### ORP1L and its association with VAPA are required for ERLAD of ATZ

Deletion of ORP1L by CRISPR/Cas9 genome editing (Fig. 5A) substantially impairs lysosomal delivery of ATZ in HeLa cells (Figs. 5B *vs*. 5C and quantifications in 5H, +pCDNA3). The lysosomal delivery of ATZ is only poorly, if at all, restored upon back transfection of ORP1L (Figs. 5D, 5H, +ORP1L-GFP). However, a partial recovery of ERLAD was obtained upon co-transfection of ORP1L and RAB7 (Figs. 5F, 5H).

**Figure 5.**
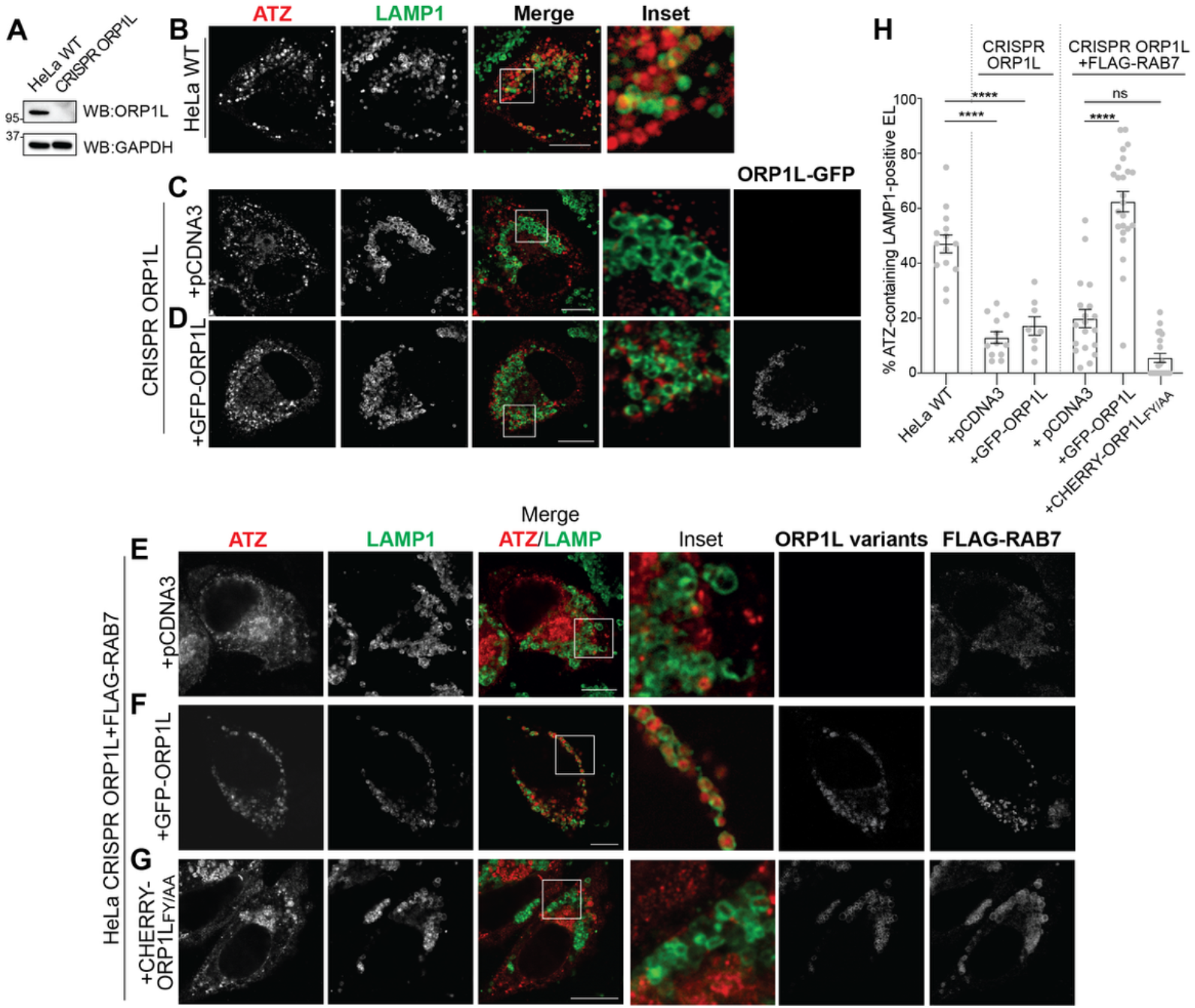
ORP1L is required for ATZ delivery to endolysosomes. (**A**) WB analysis showing the knockout of ORP1L in HeLa cells. (**B**) CLSM analysis of ATZ-HA delivery to LAMP1-positive endolysosomes in HeLa WT cells treated with 100nM BafA1. (**C**) CLSM analysis of ATZ-HA delivery to LAMP1-positive endolysosomes in HeLa CRISPR ORP1L cells mock transfected, treated with 100nM BafA1. (**D**) CLSM analysis of ATZ-HA delivery to LAMP1-positive endolysosomes in HeLa CRISPR ORP1L cells transfected with GFP-ORP1L, treated with 100nM BafA1. (**E**) CLSM analysis of HALO-ATZ delivery to LAMP1-positive endolysosomes in HeLa CRISPR ORP1L cells transfected with FLAG-Rab7, treated with 100nM BafA1. (**F**) CLSM analysis of HALO-ATZ delivery to LAMP1-positive endolysosomes in HeLa CRISPR ORP1L cells transfected with GFP-ORP1L and FLAG-Rab7, treated with 100nM BafA1. (**G**) CLSM analysis of ATZ-HA delivery to LAMP1-positive endolysosomes in HeLa CRISPR ORP1L cells transfected with CHERRY-ORP1L_FY/AA_ and FLAG-Rab7, treated with 100nM BafA1. (**H**) Quantification of **B-G** (n = 14, 12, 8, 19, 24 and 22 cells of 3 independent experiments, respectively). One-way ANOVA and Dunnett’s multiple comparisons test, ^ns^*P* > 0.05, \*\**P* < 0.01, **** *P* < 0.0001. F=56.76. Scale bar: 10μm.

The co-transfection of ORP1L_FY-AA_, a mutant form of ORP1L that does not bind VAPA^43^, with RAB7 does not rescue ATZ delivery to the degradative compartments (Figs. 5G, 5H). All in all, these data confirm that multiprotein complexes involved in VAPA:ORP1L:RAB7 contact sites contribute with the CNX:FAM134B segregation complex and the STX17:SNAP29:VAMP8 membrane fusion complex to the efficient clearance from the ER of ATZ polymers via LC3-dependent delivery to LAMP1-positive endolysosomal compartments (Fig. 6).

**Figure 6.**
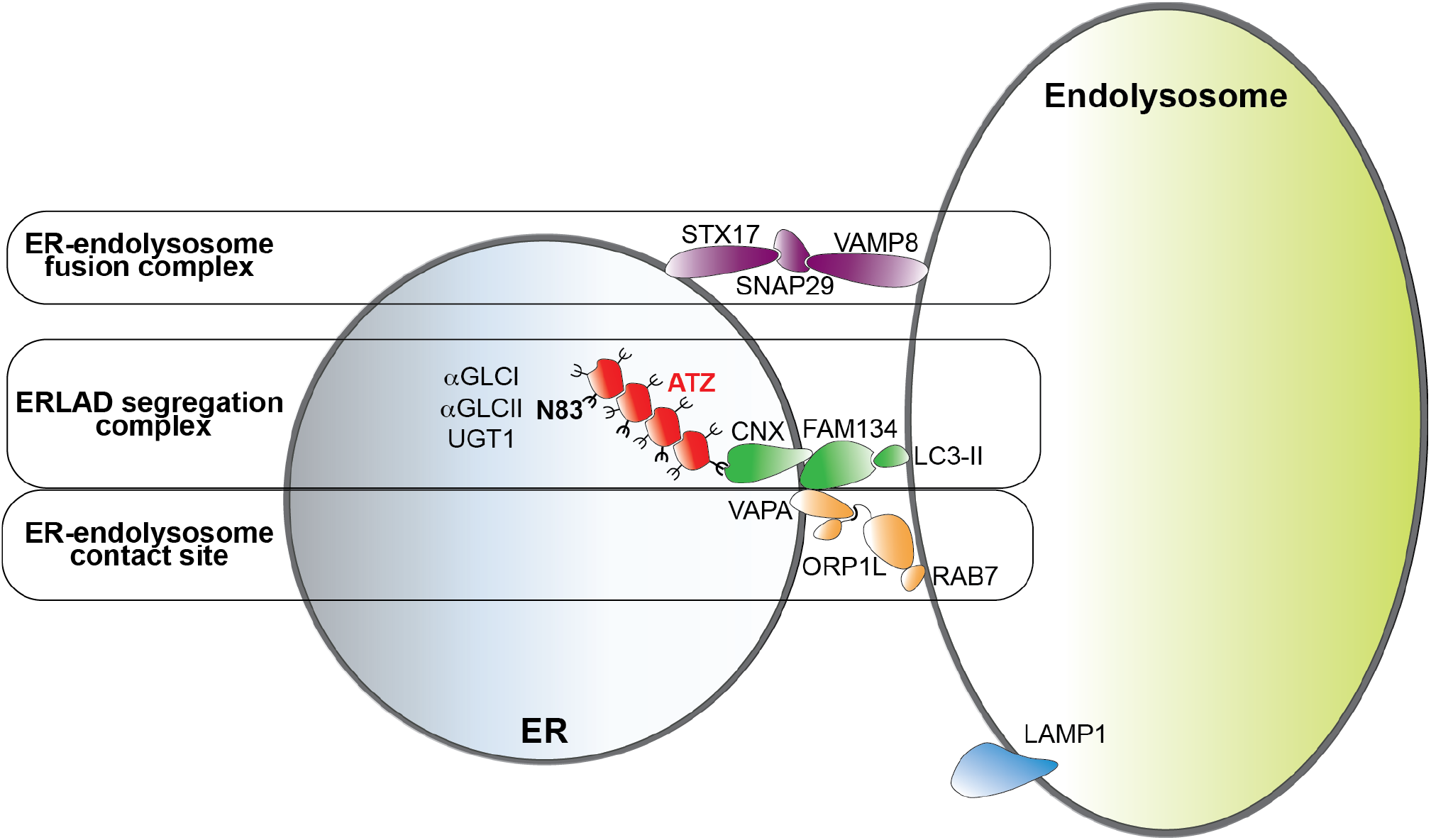
Functional complexes involved in ERLAD of ATZ polymers. Luminal expression of ATZ polymers and persistent cycles of de-glucosylation by α-glucosidase II (αGLCII) and re-glucosylation by UDP-glucose:glycoprotein glucosyltransferase 1 (UGT1) of the oligosaccharide at position 83 of the ATZ polypeptide prolong the association of ATZ polymers with the lectin chaperone calnexin (CNX)^9^. This leads to the formation of a segregation complex between CNX and the ER-phagy receptor FAM134B^10^ and, as shown here, stabilizes ER-endolysosome contacts involving the FAM134B interactor VAPA, the endolysosome-associated small GTPase RAB7 and the connecting protein ORP1L. Membrane fusion events controlled by the SNARE proteins STX17, SNAP29 and VAMP8 allow delivery of ATZ polymers within endolysosomes for clearance^10^.

## Discussion

The ER is site of protein synthesis and maturation. Sophisticated quality control pathways ensure that native and functional polypeptides are transported at their *intra-* or *extra*-cellular site of activity^44^. Terminally misfolded polypeptides are removed from the ER via proteasomal (ERAD), or lysosomal (ERLAD) pathways^1-8^. The mechanistic dissection of ERLAD is still in its infancy, and the study of the client specific pathways that deliver misfolded proteins from their site of generation (the ER) to the endolysosomal (or vacuolar in yeast and plants) degradative compartments is shedding light on intracellular quality control, client segregation, ER remodeling, transport, and clearance pathways^4^. As thoroughly reviewed elsewhere^4^, the autophagic and non-autophagic routes of misfolded proteins for lysosomal degradation include a detour via the plasma membrane (e.g., for GPI anchored misfolded polypeptides^45-47^), the capture of ER portions containing misfolded proteins by double membrane autophagosomes or by endolysosomes (e.g., for mutant forms of pro-collagen^9,48,49^), the fusion of ER portions containing misfolded proteins with degradative endolysosomes (e.g., for ATZ polymers^9,10,12^). This latter type of ERLAD, the LC3-dependent lysosomal clearance of misfolded polypeptides, relies on the persistent re-glucosylation of the misfolded polypeptide that prolongs engagement of the lectin chaperone CNX with subsequent segregation in ER subdomains displaying the ER-phagy receptor FAM134B at the limiting membrane. These ER subdomains eventually deliver their toxic content for clearance within RAB7/LAMP1-positive endolysosomes upon SNARE proteins-driven fusion^9,10,12^. Notably, genetic variants of FAM134 proteins are putative disease modifiers in children expressing ATZ^50^, revealing that this catabolic pathway, if defective, may significantly worsen disease progression and hepatic toxicity linked to the Z mutation, eventually leading to the necessity for a liver transplant. The mechanistic dissection of this pathway led to the characterization of a multimeric protein complex, whose abundance/stability is enhanced in cells expressing ATZ polymers. In this complex, the ER resident tail-anchored protein VAPA^14-21,30-33^, a major interactor of the ER-phagy receptor FAM134B at the ER membrane with unknown function in ER-phagy so far^22^ and the endolysosome-associated small GTPase RAB7 are bridged by the cytoplasmic ORP1L protein to form ER:endolysosome MCS, whose formation is required for efficient clearance of ATZ polymers from the ER. MCS between ER and endolysosomes are relevant for the ORP1L-driven transfer of lipids from the site of their biosynthesis (the ER) to endolysosomal membrane, control endolysosome maturation and, according to the results of our research, promote delivery of toxic macromolecules, in this case, disease-causing misfolded polypeptides, for removal from our cells. Notably, VAPA is a major partner of a series of ER-phagy receptors in yeast and mammalian cells^22,37,38,40,51^. For soluble ER-phagy receptors (CALCOCO1 in mammalian cells^38^ and Epr1 in yeast^40^), association with the ER proteins of the VAP family recruits the receptor at the ER membrane upon activation of the ER-phagy program. Our data imply that these associations could serve to recruit the receptor at ER subdomains in close contact with endolysosomal compartments. Significantly, VAPA has recently been identified as a major client of ER-phagy^39,41^. If VAPA and/or ER-endolysosomes contact sites established by VAPA contribute to other ER-phagy programs in addition to ERLAD, remains to be established.

## Material and Methods

### Expression plasmids

pCDNA3.1 ATZ-HA, pCDNA3.1 NHK-HA, pCDNA3.1 FAM134B-V5, pCDNA3.1 HALO-ATZ, and pCDNA3.1 HALO-NHK were prepared as previously described^10,52-55^. GFP-ORP1L was a kind gift of J. Neefjes. CHERRY-ORP1L_FY/AA_, pEF FLAG-RAB7, GFP-VAPA and GFP-VAPB were kind gifts of N. Ridgway, F. Reggiori and P. De Camilli. pCDNA5 V5-ORP1L was generated by exchanging GFP of GFP-ORP1L with V5tag.

### Antibodies

Commercial antibodies used in the study were rabbit HA (Sigma), rat HA (Novus), V5 (Thermo Fisher), GAPDH (Thermo Fisher), VAPA (Bioss), VAPB (Abcam), ORP1L (Abcam), RAB7 (Abcam), FLAG (Sigma), LAMP1 clone 1D4B (Hybridoma Bank, 1D4B was deposited to the DSHB by J.T. August), LAMP1 clone H3A4 (Hybridoma Bank, 1D4B was deposited to the DSHB by August, J.T). CNX was gift A. Helenius. Alexa-conjugated and HRP-conjugated secondary antibodies were from Thermo Fisher Scientific and Jackson ImmunoResearch, respectively.

### Cell culture, transient transfection and inhibitors

HEK293 cells were purchased from ATCC. STX17-deleted mouse embryonic fibroblasts (MEFs) were generated by CRISPR/Cas9 genome editing as previously described^10^. HeLa WT and VAPA/B DKO cells were a kind gift of P. De Camilli; HeLa CRISPR mock and CRISPR ORP1L cells were a kind gift of N. Ridgway. All cell lines were grown in DMEM supplemented with 10% FBS at 37°C, 5% CO_2_. Cells were transfected with JetPrime (Polyplus) following manufacturer’s instructions. Bafilomycin A1 (Sigma) was used 100mM for 12h or 50mM for 6h if not elsewhere stated.

### Cell lysis, immunoprecipitation and Western blot

Cells were plated on Poly-Lys (Sigma) and treated as indicated. Cells were then washed with ice cold PBS with 20mM NEM (Sigma) and lysed with 2%CHAPS (Anatrace) in HEPES buffer pH 6.8 supplemented with protease inhibitors for 20 minutes on ice.

Post-nuclear supernatants (PNS) were isolated by centrifugation at 10,600g for 10 minutes. Samples were then denatured and reduced with 2% SDS- and 100mM DTT-containing sample buffer at 95°C for 5min.

For immunoprecipitation assays, CHAPS lysates were incubated with V5-agarose beads (Sigma) at 4°C overnight. After 2 washes with 0.5%CHAPS, immunocomplexes were denatured and reduced with 6% SDS- and 10mM DTT-containing sample buffer at 95°C for 5 minutes.

Samples were then loaded on 10% Tris-glycine SDS-PAGE gels and transferred to PVDF membranes using the Trans-Blot Turbo Transfer System (Bio-Rad). Membranes were blocked with 10% Milk in TBS-T and incubated with primary and then HRP-conjugated secondary antibodies diluted in TBS-T. Membranes were then developed with WesternBright ECL or Quantum (Advansta) and signals detected with FusionFX7 VILBER (Witec). Quantifications were performed with Fiji/ImageJ.

### Confocal laser scanning microscopy

Cells were plated on Alcian blue (Sigma)-treated glasses and then transfected, treated with BafA1 and subsequently fixed with 3.7% formaldehyde in PBS at room temperature for 15 minutes. HALO-tagged proteins were labeled with TMR HaLo ligand (Promega) for 12h prior fixation.

Fixed cells were permeabilized for 15 minutes with permeabilization solution (PS, 0.05% saponin, 10% goat serum, 10 mM HEPES, 15 mM glycine). Primary antibody was diluted 1:100 with PS solution and incubated for 90 minutes. After washing for 3 times for 5 minutes, cells were then stained with Alexa-conjugated secondary antibodies for 45 minutes. Cells were rinsed with PS for 3 times and water and mounted with mount liquid anti-fade (Abberior).

Confocal images were acquired with Leica TCS SP5 or Leica Stellaris SP8 microscope with a Leica HCX PL APO lambda blue 63.0 × 1.40 OIL UV objective.

The quantifications of EL delivery per cell were executed with LysoQuant, an unbiased and automated deep learning tool for fluorescent image quantification^26^. Image processing was performed with Fiji/ImageJ and Adobe Photoshop.

### Immunogold electron microscopy

Cells were plated on alcian blue coverslips, transfected and fixed with 3.7% paraformaldehyde (PFA, Sigma). After washes in PBS and 50 mM glycine, cells were permeabilized with 0.25% saponin, 0.1% BSA, and blocked in blocking buffer (0.2% BSA, 5% goat serum, 50 mM NH_4_Cl, 0.1% saponin, 20 mM PO_4_ buffer, 150 mM NaCl). Staining with primary antibodies and nanogold-labeled secondary antibodies (Nanoprobes) was performed in blocking buffer at room temperature. Cells were re-fixed in 1% glutaraldehyde, and nanogold was enlarged with gold enhancement solution (Nanoprobes) according to the manufacturer’s instructions. Cells were post-fixed with osmium tetroxide, embedded in Epon, and processed into ultrathin slices. After contrasting with uranyl acetate and lead citrate, sections were analyzed with a Zeiss LEO 512 electron microscope. Images were acquired by a 2k bottom-mounted slow-scan Proscan camera controlled by EsivisionPro 3.2 software. Image analysis and quantification were performed with FIJI.

## Statistical analysis

Plots and statical analyses were performed with GraphPad Prism 9 (GraphPad Software Inc.). To assess the statistical significance, one-way ANOVA with Dunnett’s multiple comparisons test and unpaired two-tailed *t-*test were performed. *P*-value < 0.05 (for one-way ANOVA with Dunnett’s multiple comparisons test) or *P*-value < 0.05 (for *t*-test) were considered as statistically significant.

## Acknowledgments

We thank D. Morone and A. Raimondi for assistance in light and electron microscopy. The members of Molinari’s lab for discussions, and critical reading of the manuscript and Timothy J. Bergmann for initial experiments. We thank J. Neefjes, N. Ridgway, F. Reggiori and P. De Camilli for gifts of cell lines and plasmids.

## Funding

Swiss National Science Foundation grants 310030_214903 and 320030-227541 (MM). Alpha1-Foundation Applicakon ID: 1188512.

**Table S1.** Antibodies used. IB: Immunoblotting; CLSM: Confocal Laser Scanning Microscopy; IEM: Immuno EM

| Antibody | Manufacturer | Catalog number | Dilution or Concentration |
| --- | --- | --- | --- |
| Rat anti-LAMP1 | DSHB (J.T. August) | 1D4B | 1:50 (CLSM) |
| Mouse anti-LAMP1 | DSHB (J.T. August) | H4A3 | 1:100 (CLSM) |
| Rabbit Anti-HA | Sigma | H-6908 | 1:50 (IEM); 1:100 (CLSM); 1:1000 (IB) |
| Rat Anti-HA | Novus | NBP2-50416 | 1:1000 (CLSM) |
| Mouse Anti-V5 | Thermo Fisher Scie | R96025 | 1:100 (CLSM)/1:1000 (IB) |
| Mouse anti-V5 beads conjugated | Sigma | A-7345 | 50 µl (IP) |
| Rabbit Anti-VAPA | Bioss | Bs-4226R | 1:1000 (IB) |
| Rabbit anti-VAPB | Abcam | EPR27026 | 1:1000 (IB) |
| Rabbit anti-ORP1L | Abcam | Ab131165 | 1:1000 (IB) |
| Mouse anti-GAPDH | Millipore | MAB374 | 1:30000 (IB) |
| Rabbit anti-CALNEXIN | From A. Helenius |  | 1:1000 (IB) |
| Rabbit anti-RAB7 | Abcam | Ab-137029 | 1:1000 (IB) |
| Rabbit anti-FLAG | Sigma | F-7425 | 1:100 (CLSM) |
| Anti-mouse HRP-conjugated | Southern Biotech | 1031-05 | 1:20000 (IB) |
| Protein A HRP-conjugated | Invitrogen | 101023 | 1:20000 (IB) |
| Goat anti-rabbit AlexaFluor488-conjugated | Thermo Fisher Scie | A-21206 | 1:300 (CLSM) |
| Goat anti-mouse AlexaFluor488-conjugated | Jackson ImmunoRes | 115-545-166 | 1:300 (CLSM) |
| Goat anti-rabbit AlexaFluor568-conjugated | Thermo Fisher Scie | A-11036 | 1:300 (CLSM) |
| Goat anti-mouse AlexaFluor568-conjugated | Thermo Fisher Scie | A-11031 | 1:300 (CLSM) |
| Goat anti-rat AlexaFluor647-conjugated | Thermo Fisher Scie | A-21247 | 1:300 (CLSM) |
| Goat anti-rabbit AlexaFluor405-conjugated | Thermo Fisher Scie | A-31556 | 1:150 (CLSM) |
| Goat anti-rabbit gold-labelled | Nanoprobes | 2004 | 1:100 (IEM) |

